# Antimicrobial activity of cationic antimicrobial peptides against stationary phase bacteria

**DOI:** 10.1101/2022.08.29.505668

**Authors:** Alexandro Rodríguez-Rojas, Jens Rolff

## Abstract

Antimicrobial peptides (AMPs) are ancient antimicrobial weapons used by multicellular organisms as components of their innate immune defences. Because of the antibiotic crisis, AMPs have also become candidates for developing new drugs. Here, we show that five different AMPs of different classes are effective against non-dividing E. coli and S. aureus. By comparison, three conventional antibiotics from the main three classes of antibiotics poorly kill non-dividing bacteria at clinically relevant doses. The killing of fast-growing bacteria by AMPs is faster than that of slow-dividing bacteria and, in some cases, without any difference. Still, non-dividing bacteria are effectively killed over time. Our results point to a general property of AMPs, which might explain why selection has favoured AMPs in the evolution of metazoan immune systems. The ability to kill non-dividing cells is another reason that makes AMPs exciting candidates for drug development.

## INTRODUCTION

Antimicrobial peptides (AMPs) – small, and most of the time, cationic molecules - are crucial elements of the humoral innate immune defences of all multicellular life (Lazzaro et al., 2020). AMPs are also essential players at the host-microbiome interface (Bevins and Salzman, 2011; Mergaert, 2018). Because of their evolutionary success and diversity, AMPs are considered new antimicrobial drug candidates to alleviate the current antibiotic resistance crisis (Mookherjee et al., 2020). Currently, there are around two dozen AMPs from different origins under clinical trial (Koo and Seo, 2019).

Bacterial pathogens and bacteria, in general, encode several conserved and essential genes in their genomes, where inhibition could lead to bacterial growth arrest or killing. These genes are usually the targets of all known antibiotics. Such druggable pathways range from several tens to hundreds of genes (Juhas et al., 2011). It is striking, however, that metazoan immune effectors do not exploit these easy targets while chemical defences of microbes do to gain competitive advantages (Letten et al., 2021). One possibility is that resistance evolution against antibiotics is relatively easy and ubiquitous (Blázquez et al., 2018). Toxicity or microbiome damage could be another reason to avoid using chemicals such as antibiotics for our chemical defences (Blaser, 2016).

We have been studying why AMPs were selected during evolution and what properties made them more suitable as an antimicrobial defence strategy of metazoan than other types of molecules, such as antibiotics. These insights have also the potential to inform the application of AMPs as drugs. For example, we have found that AMPs differ significantly from conventional antibiotics: including their pharmacodynamics, resulting in narrower mutant selection windows (Yu et al., 2018). In contrast to conventional antibiotics, AMPs do not increase the mutation rate even at sub-lethal concentrations (Rodríguez-Rojas et al., 2014, 2015), and they do not increase recombination frequency (Rodríguez-Rojas et al., 2018). Taken together these features of AMPs combine to lower the probabilities of resistance evolution (Yu et al., 2018). It seems that the emergence of AMPs as an antimicrobial weapon during evolution depends not on a single feature but several.

There are many physiological or pathological situations where microbes are slow growing, andthe host needs to control them. For instance, *Escherichia coli*, a typical colonizer of humans and warm-blooded animals (Martinson and Walk, 2020) grows very fast in laboratory conditions, with a doubling time or generation time around 20 minutes. The proliferation rate in human guts, however, was estimated to be near 40 hours (Savageau, 2015). Situations such as this motivated our study.

We note that slow growing bacteria also occur in biofilms, a situation we do not study here. It has been shown that some antimicrobial peptides have anti-biofilm activity (Huan et al., 2020). However, it should be noticed that many of them are different from the classical AMPs from multicellular organism. The positive charges although being essential for antimicrobial action also represent an impediment to natural AMPs penetration into biofilms (Yasir et al., 2018). The main reason lies in the fact that polysaccharides (EPS) of the biofilm matrix is negatively charged and can trap positive AMPs. This undermines the activity of AMPs. Moreover, many AMPs in biofilms are subject to hydrolytic and proteolytic breakdown (Galdiero et al., 2019).

**Stationary phase** bacteria are significantly less susceptible to antibiotics than fast-growing counterparts (Gutierrez et al., 2017; Mccall et al., 2019). Conventional antibiotics are most often only effective when bacteria are actively dividing (Eng et al., 1991; Lobritz et al., 2015). It is known that colistin, a cationic antimicrobial peptide of microbial origin (Biswas et al., 2012), kills bacteria regardless of their metabolic state (Singhal et al., 2022). Based on a similar mode of action that colistin shares with other antimicrobial peptides, we hypothesise that the ability to kill **stationary phase** bacteria is a general property of cationic AMPs from multicellular organisms. We specifically investigate whether AMPs can kill non-dividing bacteria and how the bacterial physiology of the stationary phase change this dynamic. We use five antimicrobial peptides from different origins (cecropin A, indolicidin, LL-37, melittin, and pexiganan) that are well characterised in their activities. For comparison, we also use three bactericidal drugs representing the three most relevant antibiotic families: beta-lactams, aminoglycosides and fluoroquinolones (ampicillin, gentamycin and ciprofloxacin, respectively). To extend the validity of our research, we carried out experiments using a Gram-negative bacterium (*Escherichia coli*) and a Gram-positive bacterial model (*Staphylococcus aureus*).

## RESULTS AND DISCUSSION

First, we determined the minimal inhibitory concentration (MIC) for all antimicrobials (Table 1) and used these values as a reference for the killing experiments. Then, we exposed equivalent numbers of both bacterial species to a ten-fold concentration (10X MIC) of cecropin A, indolicidin, LL-37, melittin, and pexiganan. We also used the same treatments with ampicillin, ciprofloxacin and gentamicin. Finally, we measured bacterial survival at two-time points, 4 and 24 hours post exposure for actively dividing bacteria (exponential phase) and slow-replicating ones (stationary phase). The results from these experiments are shown in Fig. 1 and Fig. 2 for *E. coli* and *S. aureus*, respectively. **In this article, we refer to fast or actively replicating bacteria as the state of the bacterial population at the time of the addition of antimicrobials**. In our experimental conditions, all antimicrobials drastically reduced bacterial counts at 4 and 24 hours of treatment for the exponentially growing bacteria. For stationary phase bacteria, all AMPs reduced viability. However, after 4 hours of exposure, there was some delay in killing by cecropin A, LL-37, melittin, and pexiganan while killing stationary phase bacteria compared to exponentially growing microbes. Indolicidin was the most efficient AMP, capable of completely killing (or reducing bacterial counts below the detection limit), bacterial cultures from both species but also for fast and slow-replicating bacteria for both time-points, 4 and 24 hours.

**Table 1.** Minimal inhibitory concentration (MIC) values for *E. coli* K12 and *S. aureus* SH1000 for different antimicrobials used in this study.

| Antimicrobials | Class | MIC (µg/ml) |  |
| --- | --- | --- | --- |
|  |  | *E. coli* K12 | *S. aureus* SH1000 |
| ampicillin | beta-lactam | 2 | 4 |
| ciprofloxacin | fluoroquinolone | 0.125 | 0.062 |
| gentamicin | aminoglycoside | 1 | 4 |
| cecropin A | cationionic AMP | 4 | 8 |
| indolicidin | cationionic AMP | 16 | 8 |
| LL-37 | cationionic AMP | 4 | 8 |
| melittin | cationionic AMP | 2 | 2 |
| pexiganan | cationionic AMP | 4 | 8 |

**Fig. 1.**
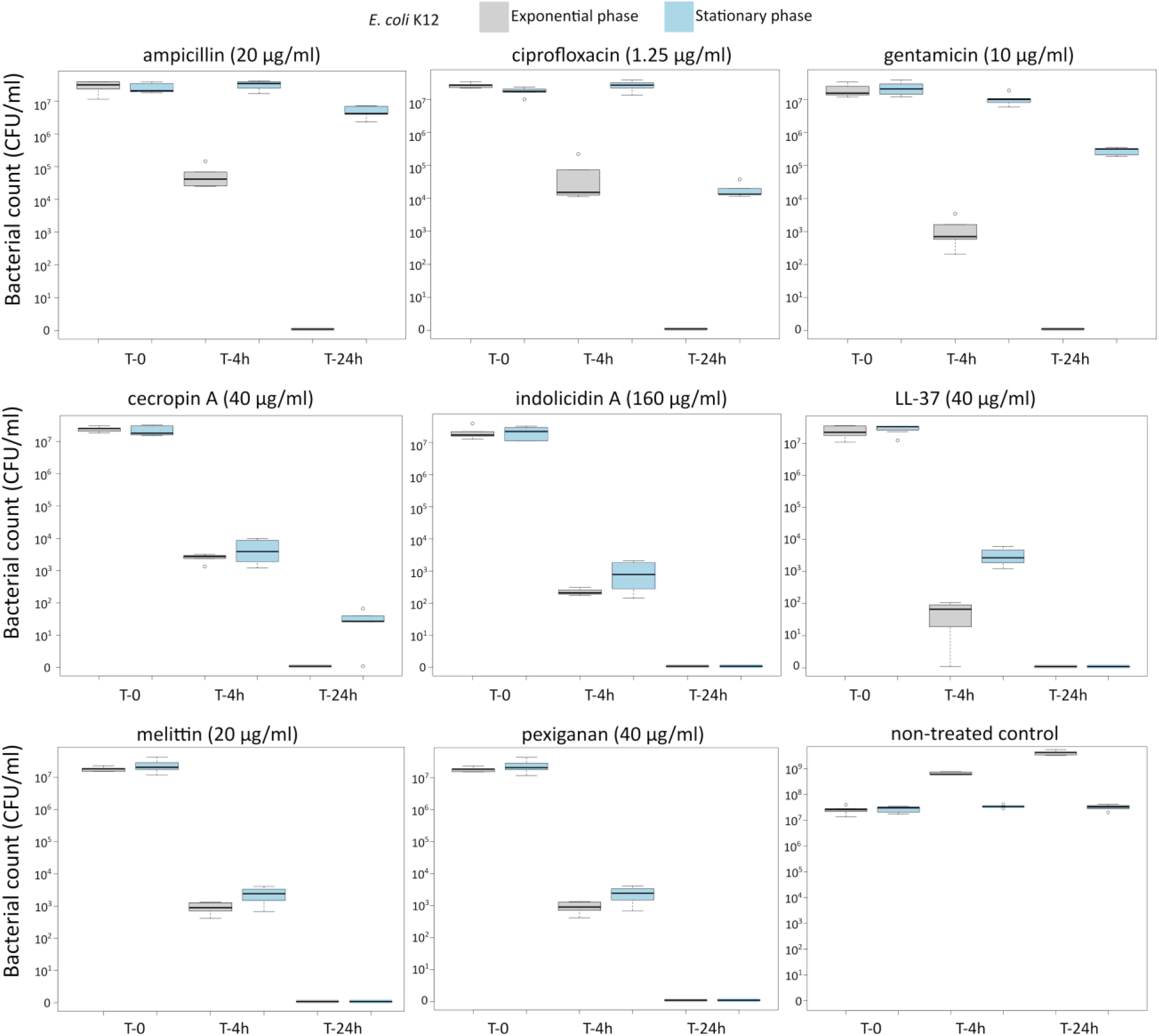
Killing of *E. coli* K12 at 10X MIC for five antimicrobial peptides and three different antibiotics. The grey boxes indicate bacteria growing exponentially while the blue ones denote bacteria treated in the stationary phase in spent medium prepared as described in the Material and Methods section.

**Fig. 2.**
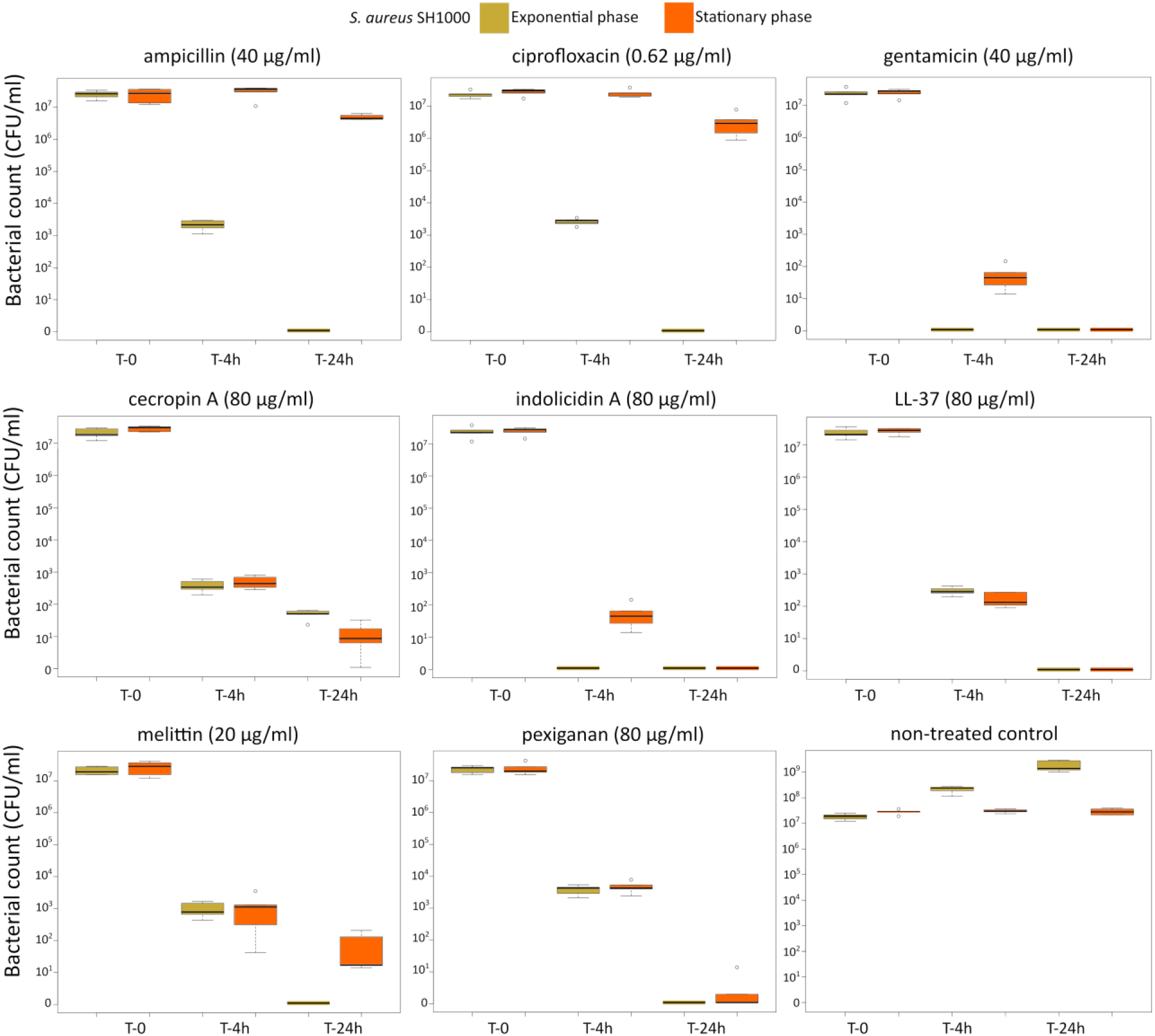
Killing of *S. aureus* SH1000 at 10X MIC for five antimicrobial peptides and three different antibiotics. The yellow light green boxes indicate bacteria growing exponentially while the orange ones denote treated bacteria in the stationary phase in spent medium prepared as described in the Material and Methods section.

The three antibiotics were very efficient in killing after 4 hours for both bacterial species of exponentially growing cells while failing to kill **stationary phase** ones at 4 hours. In particular, ampicillin did not kill stationary phase bacteria 4 hrs post treatment, with a poor killing capacity even at 24 hours. Also after 24 hours of exposure, gentamicin and ciprofloxacin reduced microbial count for both models with a significantly lower efficacy than killing fast-replicating cultures. All statistical inferences of the killing rate at 4 and 24 hours for each antimicrobial and both bacterial species are provided in Table 2 and Table 3.

**Table 2.** Killing fold-change of *E. coli* K12 growing exponentially versus stationary phase cultures and their comparative statistical difference (Welsh’s test).

| Antimicrobials | *E. coli* K12 |  |  |  |
| --- | --- | --- | --- | --- |
|  | Killing fold-change 4 hours (median) | *p*-value | Killing fold-change 24 hours | *p*-value |
| ampicillin | 595.18 | 0.0183 | 4150000 | 0.0033 |
| ciprofloxacin | 1072 | 0.0018 | 13300 | 0.0085 |
| gentamicin | 10380.62 | 0.0041 | 317000 | 0.0006 |
| cecropin A | 1.21 | ns | 27 | 0.0220 |
| indolicidin | 9.09E-02 | 0.0027 | 1 | ns |
| LL-37 | 3.71 | 0.0235 | 1 | ns |
| melittin | 65.79 | 0.0007 | 1 | ns |
| pexiganan | 2.53 | 0.0168 | 1 | ns |

**Table 3.**
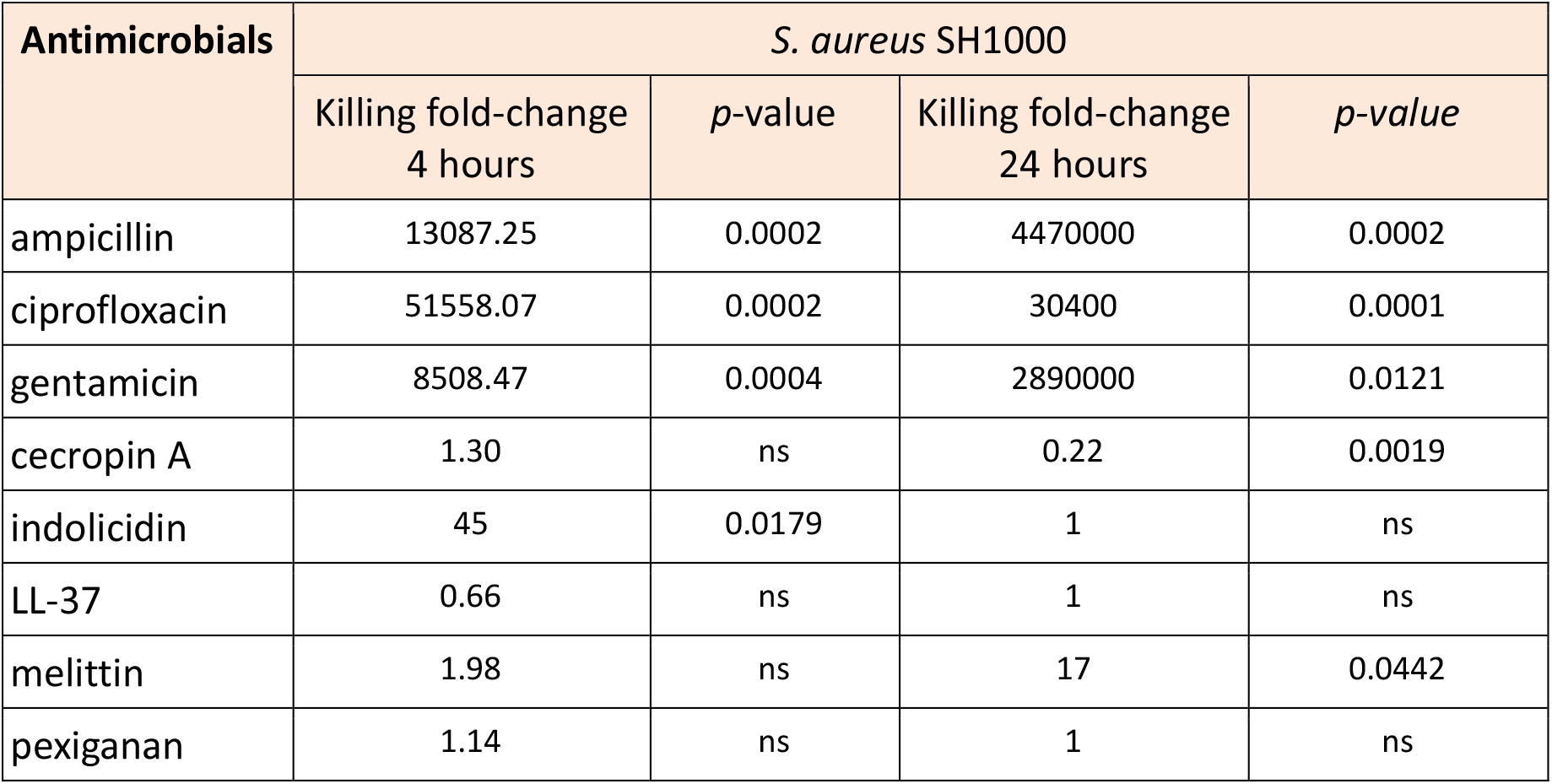
Killing fold-change of *S. aureus* SH1000 growing exponentially versus stationary phase cultures and their comparative statistical difference (Welsh’s test).

Indolicidin, the most effective AMP at killing non-dividing bacteria, is a 13-residue peptide belonging to the cathelicidin family, with a broad-spectrum activity against a wide range of targets, such as Gram-positive and Gram-negative bacteria, fungi and viruses (Batista Araujo et al., 2022). This peptide was isolated from neutrophil blood cells of cows (Selsted et al., 1992). Our findings suggest that the capacity of cationic AMPs to kill slow-replicating bacteria is rather common and may be conserved across the tree of life. In this work, we also used insect derivative peptides such as cecropin A, first isolated from the hemolymph of the moth *Hyalophora cecropia* (Lee and Brey, 1994), and melittin from the venom of honeybees (Habermann and Jentsch, 1966). In addition to indolicidin, we used two more vertebrate AMPs, pexiganan, a derivative of magainin II from the skin of the African frog *Xenopus laevis* (Ge et al., 1999) and the human peptide LL-37, which also has antibiofilm activity (Ridyard and Overhage, 2021). All these five AMPs were efficient at killing non-dividing (stationary phase) bacteria. This property adds to the potential benefits of AMPs as antimicrobial drug candidates but also to our understanding of their role and evolution as main components of metazoan innate immune defenses.

All antimicrobials are sensitive to the inoculum effect or cell density, a phenomenon that decreases their efficacy (Udekwu et al., 2009). This is one of the reasons why for example, biofilms are less sensitive to antimicrobials in general. This is particularly true for positively charged drugs, mostly aminoglycosides and the cationic antimicrobial peptide colistin with a very drastic diminishing of the killing activity within biofilm microenvironments (Kirby et al., 2012). The lower killing of AMPs due to high density bacterial population seems to be a prevalent phenomenon (Loffredo et al., 2021). Although these problems have hampered the utilization of AMPs as drugs, efforts to make AMPs more stable by chemical modifications are widespread and could help to mitigate this issue. Because of the inoculum effect, we designed this study to investigate bacterial killing in low-density populations, a situation, which we think, is common enough to warrant study. During the onset of an infection, it is common that microbes enter the body in low proliferating state or close to the stationary phase. They usually are also in small numbers, where AMPs would work well as a first line of defence. In the same line, it is also known that the inoculum size plays a fundamental role in the probability of establishing an infection (Grant et al., 2008). Finally, we would like to mention that a possible limitation of this study is that antimicrobial peptides could behave differently within the host compared to *in vitro* conditions. Therefore, future studies will be necessary to study how the cationic antimicrobial peptides kill stationary phase or slow replicating bacteria *in vivo*.

## MATERIAL AND METHODS

### Bacteria and growth conditions

The *Escherichia coli* K12 from Yale University Microbial Stock Center and *Staphylococcus aureus* SH1000 (Horsburgh et al., 2002) were used for all experiments. All cultures related to antimicrobial tests and experiments were carried out in Mueller-Hinton I Broth (Sigma) devoid of cations. Both strains were routinely cultured in Lysogeny Broth (LB medium).

### Antimicrobials

For this study, we used five different AMPs (cecropin A, indolicidin, LL-37, melittin, and pexiganan) and three antibiotics (ampicillin, gentamycin and ciprofloxacin, respectively). All antimicrobials were purchased from Sigma except pexiganan that was a generous gift from Dr. Michael A. Zasloff from Georgetown University.

### Minimal inhibitory concentration (MIC)

The Minimal inhibitory concentration for each antimicrobial were determined according to CLSI recommendations by the microdilution method (CLSI, 2018) with minor modifications for antimicrobial peptides (Giacometti et al., 2000). For comparison, we kept these modifications also for antibiotics. Inoculum size that was adjusted to approximately 1×10^6^ CFU/ml from a 2-hour mid-exponential phase obtained by diluting 100 μl of overnight cultures in 10 ml of fresh medium in 50 ml Falcon tubes). The MIC was defined as the minimal antimicrobial concentration that inhibited growth, after 24 h of incubation in liquid MHB medium at 37°C. Polypropylene non-binding plates (96 wells, Th. Geyer, Germany) were used for all experiments. Results are presented in Table 1.

### Fast-growing bacteria killing experiment

For exponentially growing bacteria, five independent cultures per treatment were diluted 1:100 from a 16-hour overnight culture (by adding 100 μl to 10 ml of MHB medium in 50 ml Falcon tubes). Then, the bacteria were grown for 2 hours for *E. coli* and 2.5 hours for *S. aureus*, to reach approximately 2 × 10^8^ CFU/ml. The cultures were then diluted 1/100 in fresh medium to reach approximately 2 × 10^6^ CFU/ml. Two ml of diluted culture were exposed to 10x MIC to the AMPs and the antibiotics. Volumes of 100 μl-sample were extracted at 4 and 24 hours. The aliquots were diluted and plated to determine cell viability. Non-treated cells were used as a control. All incubations took place at 37°C with shaking.

### Generation of a stationary phase-like culture medium (spent medium)

In order to maintain bacteria in their non-dividing state, we generated a medium to carry out the killing assays. From the supernatant of 500 ml of 48-hour culture from each bacterium, the cells were pelleted at 4 000 x g for 30 minutes. The pH of each culture was adjusted to match the original pH of 7.2. To remove additional cell debris, flagella rest and outer membrane vesicles that could potentially interfere with the activity of AMPs (Manning and Kuehn, 2011), the supernatants were ultracentrifuged at 100 000 x g during 16 hours at +4 °C. Thereafter, 250 ml of the supernatant from each flask was carefully recovered, without perturbing the pellet, and aseptically filtrated using 0.22 μm syringe filters and 40 ml were transferred to sterile 50 ml-falcon tubes, that were stored at -20 °C until use. We refer to this medium as spent medium. The medium was tested to show its incapacity to sustain additional bacterial growth.

### Non-dividing bacteria killing experiment

For stationary phase bacteria, five independent cultures per treatment were used. Bacteria from 48-hour cultures containing approximately 3 × 10^9^ CFU/ml for *E. coli* and 2 × 10^9^ CFU/ml for *S. aureus* were diluted in spent medium (see preparation above) to a final cell density of 2×10^6^ CFU/ml. Then, the bacteria were treated identically to the *fast-growing bacteria killing experiment* described above.

### Statistical analysis

Statistical testing and plots were done in R version 3.3.2 (R Core Team, 2017), using Rstudio version 1.0.143 (R Development Core Team, 2015). To compare killing rates between bacteria growing in exponential and stationary phase for each antimicrobial after 4 and 24 hours of exposure, a Welch’s t-test was used. Values below the detection limit (zero colony counts) were imputed for statistical purposes assigning a value of 1.

## Acknowledgments

We would like to thank Dr. Michael A. Zasloff from Georgetown University for kindly providing pexiganan and Greta Santi from Padova University for initial technical assistance.

## Data Availability Statement

All relevant data are within the manuscript and its supporting Information files, including raw data are available with the manuscript.

